# Umap and Bismap: quantifying genome and methylome mappability

**DOI:** 10.1101/095463

**Authors:** Mehran Karimzadeh, Carl Ernst, Anshul Kundaje, Michael M. Hoffman

## Abstract

**Motivation:** Short-read sequencing enables assessment of genetic and biochemical traits of individual genomic regions, such as the location of genetic variation, protein binding, and chemical modifications. Every region in a genome assembly has a property called *mappability* which measures the extent to which it can be uniquely mapped by sequence reads. In regions of lower mappability, estimates of genomic and epigenomic characteristics from sequencing assays are less reliable. At best, sequencing assays will produce misleadingly low numbers of reads in these regions. At worst, these regions have increased susceptibility to spurious mapping from reads from other regions of the genome with sequencing errors or unexpected genetic variation. Bisulfite sequencing approaches used to identify DNA methylation exacerbate these problems by introducing large numbers of reads that map to multiple regions. While many tools consider mappability during the read mapping process, subsequent analysis often loses this information. Both to correct assumptions of uniformity in downstream analysis, and to identify regions where the analysis is less reliable, it is necessary to know the mappability of both ordinary and bisulfite-converted genomes.

**Results:** We introduce the Umap software for identifying uniquely mappable regions of any genome. Its Bismap extension identifies mappability of the bisulfite-converted genome. With a read length of 24 bp, 18.7% of the unmodified genome and 33.5% of the bisulfite-converted genome is not uniquely mappable. This complicates interpretation of functional genomics experiments using short-read sequencing, especially in regulatory regions. For example, 81% of human CpG islands overlap with regions that are not uniquely mappable. Similarly, in some ENCODE ChIP-seq datasets, up to 50% of peaks overlap with regions that are not uniquely mappable. We also explored differentially methylated regions from a case-control study and identified regions that were not uniquely mappable. In the widely used 450K methylation array, 4,230 probes are not uniquely mappable. Genome mappability is higher with longer sequencing reads, but most publicly available ChIP-seq and reduced representation bisulfite sequencing datasets have shorter reads. Therefore, uneven and low mappability remains a concern in a majority of existing data.

**Availability:** A Umap and Bismap track hub for human genome assemblies GRCh37/hg19 and GRCh38/hg38, and mouse assemblies GRCm37/mm9 and GRCm38/mm10 is available at http://bismap.hoffmanlab.org for use with the UCSC and Ensembl genome browsers. We have deposited in Zenodo the current version of our software (https://doi.org/10.5281/zenodo.800648) and the mappability data used in this project (https://doi.org/10.5281/zenodo.800645). In addition, the software (https://bitbucket.org/hoffmanlab/umap) is freely available under the GNU General Public License, version 3 (GPLv3).

## 1 Introduction

High-throughput sequencing enables low-cost collection of high numbers of sequencing reads but these reads are often short. Short-read sequencing limits the fraction of the genome that we can unambiguously sequence by aligning the reads to the reference genome (Figure 1b). Still, we can identify much of the regulatory regions of the genome such as transcription factor binding sites, histone modifications and other important regulatory regions. However, reads that are ambiguously mapped produce a false positive signal that misleads analysis. Some regions of the genome with low complexity including repeat elements are not uniquely mappable at a given read length. Other regions overlap few uniquely mappable reads, and consequently the mappability is low. To map the regions with low mappability, a high sequencing depth is required to assure that sequencing reads completely overlap with few uniquely mappable reads in that region. If sequencing depth is low and genomic variation or sequencing error is high, the signal from a low mappability region is biased by reads falsely mapped to that region.

**Figure 1:**
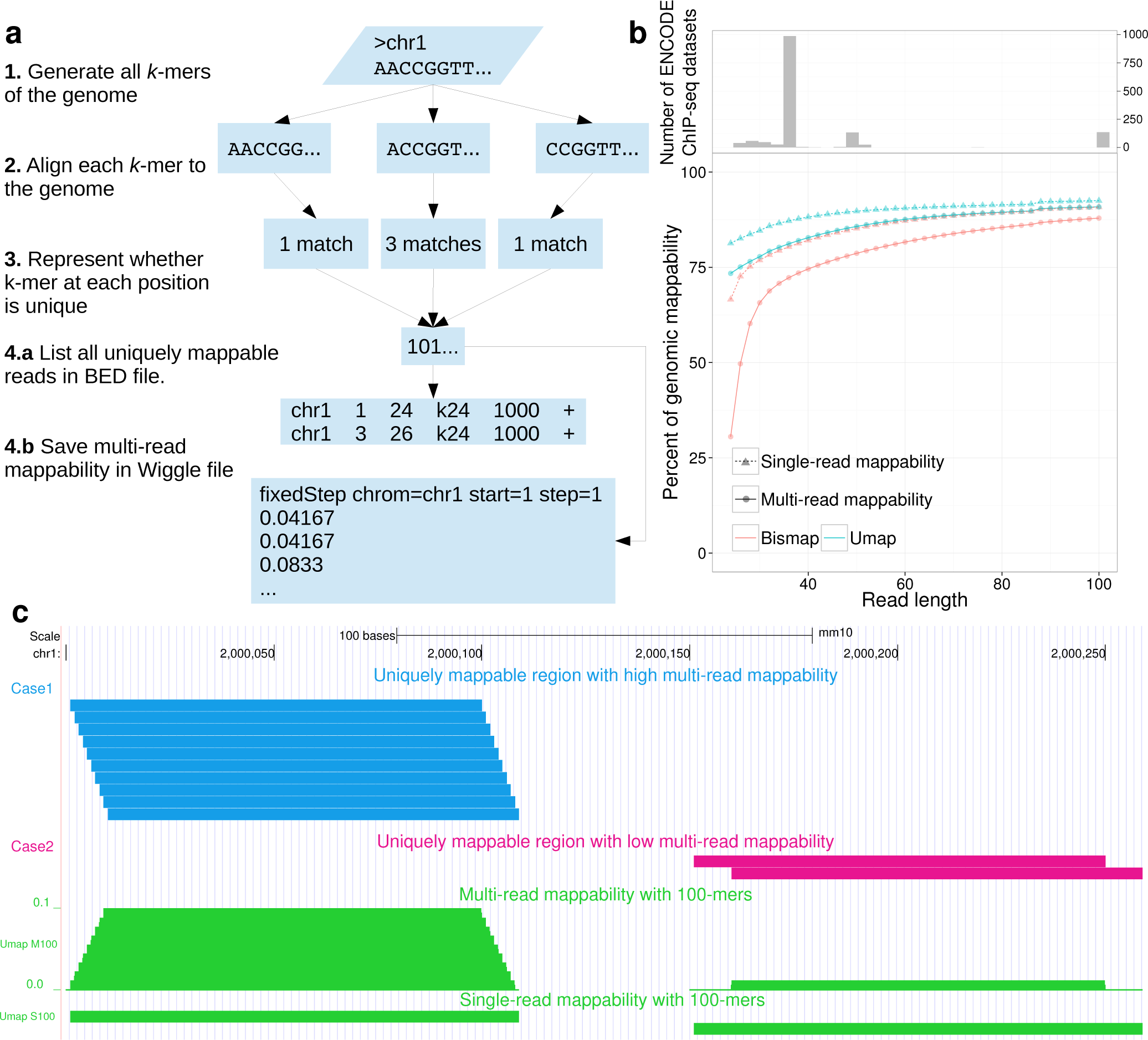
Mappability of the genome by Umap. (**a**) The Umap workflow identifies all unique k-mers of a genome given a read length of k. (**b**) Mappability of the human genome and methylome for read lengths between 24 and 100. (**c**) All of the uniquely mappable reads in two regions with high and low multi-read mappability is shown. In *Case 1* (blue), all possible reads covering the region are uniquely mappable. In *Case 2* (magenta), only two reads out of 10 are uniquely mappable.

Most short-read alignment algorithms determine if any read maps to one or more regions in the genome. However, one must consider this in context of the surrounding regions, even if a read maps uniquely. A single nucleotide change might change a read from uniquely mappable to not. A uniquely mappable read that aligns to a region with low mappability, has a high chance of mapping incorrectly due to genetic variation or sequencing error.

In bisulfite sequencing, this problem increases. Bisulfite treatment reduces unmethylated cytosine to uracil (sequenced as **T**) while 5-methylcytosine remains intact (sequenced as **C**). Bisulfite treatment significantly increases the number of repeated short sequences in the genome. Many regions uniquely mappable in an unmodified genome no longer uniquely map after bisulfite conversion. Incorrect mapping of bisulfite sequencing reads creates a false methylation signal that can bias downstream analysis and interpretation. When confounding factors such as read length, sequencing depth or mutation rate differ among cases, this bias becomes even more evident.

In an unmodified human genome, 18.7% of the 24-mers do not map uniquely (Figure 1b). This quantity increases to 33.5% for a bisulfite-converted genome (Figure 1b). In certain cases, the difference between a uniquely mappable and a non-uniquely mappable read can be only one nucleotide. Sequencer base-calling errors and genetic variation often affect alignment, but we cannot comprehensively account for them. These biases further exacerbate alignment when the read length is shorter, emphasizing the importance of considering genomic mappability in any analysis involving short-read sequencing. While previous tools such as the GEM mappability software^1^ identify mappability of the genome, no existing software solves the methylome mappability problem. In addition, existing tools prove difficult to use or lack available source code. To solve this problem, we developed the Umap software, with a bisulfite mappability extension called Bismap.

## 2 Methods

### 2.1 Single and multi-read mappability

Umap identifies the uniquely mappable reads of any genome for a range of sequencing read lengths. The Bismap extension of Umap produces uniquely mappable reads of a bisulfite-converted genome. Both Umap and Bismap produce an integer vector for each chromosome that defines the mappability for any region and can be converted to a browser extensible data (BED) file. One way to assess mappability of a genomic region is by the **single-read mappability** — the fraction of that region which overlaps with at least one uniquely mappable *k*-mer.

Analysis of sequencing data involves inferences about a base’s genetic or regulatory state from observations of all reads overlapping that base. Therefore, we must consider the mappability of all reads overlapping a position or region, when estimating how many mapped reads we might expect. Single-read mappability assumes that uniquely mappable reads are uniformly distributed in the genome, while in reality we observe frequent localized enrichment of uniquely mappable reads.

A region can have 100% single-read mappability, but a below-average number of uniquely mappable reads that can overlap that region (Figure 1c). For example, a 1 kbp region with 100% single-read mappability can be mappable due to a minimum of 10 unique non-overlapping 100-mers or a maximum of 1100 unique highly overlapping 100-mers. Therefore, we define the **multi-read mappability** — the probability that a randomly selected *k*-mer in a given region is uniquely mappable. For the genomic region *G_i_*_:*j*_ starting at *i* and ending at *j*, there are *j — i* + *k* + 1 different *k*-mers that overlap with *G_i:j_*. The multi-read mappability of *G*_i:j_ is the fraction of those *k*-mers that are uniquely mappable (Figure 1c).

### 2.2 Mappability of the unmodified genome

Umap uses three steps to identify the mappability of a genome for a given read length *k* (Figure 1a). First, it generates all possible *k*-mers of the genome. Second, it maps these unique *k*-mers to the genome with Bowtie^2^ version 1.1.0. Third, Umap marks the start position of each *k*-mer that aligns to only one region in the genome. Umap repeats these steps for a range of different *k*-mers and stores the data of each chromosome in a binary vector *X* with the same length as the chromosome’s sequence. For read length *k*, *X_i_* = 1 means that the sequence starting at *X_i_* and ending at *X_i+k_* is uniquely mappable on the + strand. Since we align to both strands of the genome, the reverse complement of this same sequence starting at *X_i+k_* in the — strand is also uniquely mappable. *X_i_* = 0 means that the sequence starting at *X_i_* and ending at *X_i+k_* can be mapped to at least two different regions in the genome.

Eventually, Umap merges data of several read lengths to make a compact integer vector for each chromosome (Figure 1a, step 3). In this vector, non-zero values at position *X_i_* indicate the smallest *k*-mer that position *X_i_* to *X_i+K_* is uniquely mappable with, where *K* is the largest *k*-mer in the range. For example *X_i_* = 24 means that the region *X_i_* to *X_i_*_+24_ is uniquely mappable. This also means that any read longer than 24 nucleotides that starts at *X_i_* is also uniquely mappable.

Umap translates these integer vectors into six-column BED files for the whole genome (Figure 1a, step 4). Additionally, Umap can calculate single-read mappability and multi-read mappability for specified regions in any input BED file.

Although Bowtie can align with mismatches, here we do not use this capability. By defining mappability with exact matches only, we provide baseline identification of regions that are not uniquely mappable no matter how high the sequencing coverage. Nonetheless, the Umap software allows users to change alignment options, including mismatch parameters.

### 2.3 Mappability of the bisulfite-converted genome

To identify the single-read mappability of a bisulfite-converted genome, we create two altered genome sequences (Figure 2). In the first sequence, we convert all cytosines to thymine (**C**→**T**). In the other sequence we convert all guanines to adenine (**G**→**A**). Our approach follows those of Bismark^3^ and BWA-meth^4^. We convert the genome sequence this way because bisulfite treatment converts un-methylated cytosine to uracil which is read as thymine. Similarly the guanine that is base-pairing with the un-methylated cytosine in the — strand converts to adenine. These two conversions, however, never occur at the same time on the same read. We identify the uniquely mappable regions of these two genomes separately, and then combine the data to represent the single-read mappability of the + and — strands in the bisulfite-converted genome. For an unmodified genome, however, the mappability of the + and — strand is identical by definition.

**Figure 2:**
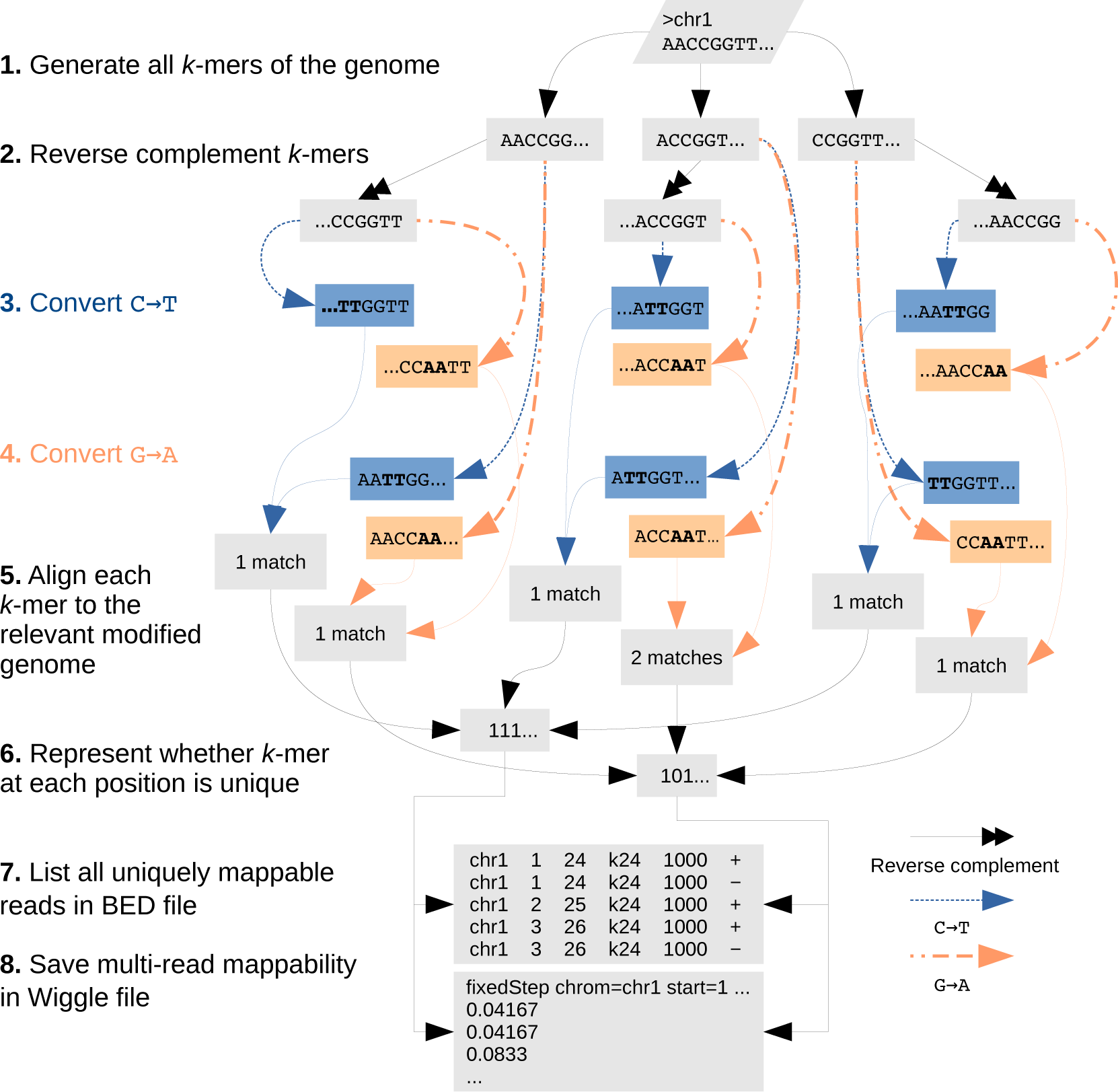
Mappability of the methylome by Bismap. Bismap identifies uniquely mappable k-mers of a bisulfite-converted genome. It simulates the same changes that may occur in bisulfite treatment on the + strand (`C→T`) and — strand (`G→A`). To account for sequence of the — strand, we generate an extra set of reverse-complemented chromosomes and then simulate bisulfite conversion on these chromosomes. We don’t simulate reverse complementation after bisulfite conversion, because the experimental protocol does not involve post-conversion DNA amplification. We then align *k*-mers by disabling complement search and combine the resulting data to quantify the mappability of a bisulfite-converted genome.

Bismap requires special handling of reverse complementation of `C→T` or `G→A` converted genomes. Conversion of `C→T` on the sequence 5'— AATTCCGG —3' produces 5'— AATTTTGG —3'. In the Bowtie index, the reverse complement of the latter would be 5'— CCAAAATT —3'. For the purpose of identifying the mappability of the bisulfite-converted genome, however, we expect the reverse complement to be derived from the original converted sequence, yielding 5' — CCGGAATT —3', and then after `C→T` conversion, 5'— TTGGAATT —3'. Both + and — strands undergo bisulfite treatment simultaneously, and there is no DNA replication to create new reverse complements after bisulfite treatment. To handle this issue, Bismap creates its own reverse complemented chromosomes and suppresses Bowtie’s usual reverse complement mapping.

Umap and Bismap each take ~ 200 core-hours on a 2.6 GHz Intel(R) Xeon CPU E5-2650 v2 processor and less than 500 MB of memory to run for some read length. This is a massively parallelizable task, so on a computing cluster with 400 cores, the task takes only 30 min of wall-clock time.

### 2.4 ENCODE ChIP-seq experiments

We downloaded ENCODE^5^ chromatin immunoprecipitation-sequencing (ChIP-seq) FASTQ files from the ENCODE Data Coordination Center^6^ and aligned them to GRCh38 using Bowtie^7^ 2. We switched to Bowtie 2 for this analysis because it supports gapped alignment, which we didn’t need for mappability calculations.

We used Samtools^8^ to remove duplicated sequences and those with a mapping quality of < 10. This assures that the probability of correct mapping to the genome for any read is > 0.9. Pooling replicates from the same experiment, we used MACS^9^ version 2 with —`nomodel` and —`qvalue 0.001` options to identify ChIP-seq peaks. Finally, Umap measured single-read mappability and multi-read mappability within the peaks.

### 2.5 CpG islands

We downloaded CpG islands^10^ for GRCh38 from the UCSC Genome Browser^11^ (http://epigraph.mpi-inf.mpg.de/download/CpG_islands_revisited). These CpG islands come from a hidden Markov model (HMM) fitted to genomic G+C content. We then annotated CpG features around the CpG islands following published definitions^10,12^ (Table 1). Then we used Umap and Bismap to measure mappability across these annotations.

**Table 1:** CpG annotations.

| Annotation | Definition |
| --- | --- |
| CpG island | HMM fitted to G+C content |
| CpG shore | 2 kbp area surrounding CpG islands |
| CpG shelf | 2 kbp area surrounding CpG shores |
| CpG resort | Collection of islands, shores and shelves |

### 2.6 Whole-genome bisulfite sequencing analysis

First, we obtained datasets of whole-genome bisulfite sequencing of murine mammary tissues^13^ from the Sequence Read Archive (accession numbers SRR1946823, SRR1946824, SRR1946819, and SRR1946820). Second, we trimmed Illumina TruSeq adapters from FASTQ files with Trim Galore^14^. Third, for each experiment, we break down sequencing reads to produce two different FASTQ files with read lengths of 50 bp and 100 bp. For example, if the read length of an experiment is 182 bp and we want to generate a FASTQ file with read length of 50 bp, each sequencing read would produce three different 50-bp sequencing reads (we would not use the remaining 32bp). We aligned these modified FASTQ files with BWA-meth ^4^ to the GRCm38 genome. We extracted CpG-context methylation using PileOmeth^15^. We use BSmooth ^16^ (version 0.4.2) for identifying differentially methylated regions. Finally, we used Bismap to measure mappability of differentially methylated regions with at least four CpG dinucleotides.

### 2.7 Other methylation assays

DiseaseMeth^17^, a human methylation database, provides access to 17,024 methylation datasets from 88 different human diseases. These data are a collection of experiments using various platforms, including 2,728 assays using the Illumina Infinium HumanMethylation27 (27K) BeadChip, and 9,795 assays using the Illumina Infinium HumanMethylation450 (450K) BeadChip. To identify which 50 bp probe sequences ^18,19^ do not map uniquely to the GRCh37 genome, we measured single-read mappability with Umap. To identify which probes do not map uniquely after bisulfite conversion, we measured single-read and multi-read mappability with Bismap.

In addition, we examined whether the exact 50-mer probe sequence mapped uniquely.

DiseaseMeth also contains 71 experimental datasets using reduced representation bisulfite sequencing (RRBS)^20^. For CpG dinucleotides captured in RRBS experiments and annotated by DiseaseMeth, we examined the multi-read mappability for read lengths of 24 bp, 36 bp, 50 bp, and 100 bp.

### 2.8 Umap and Bismap track hub

We used read lengths of 24 bp, 36 bp, 50 bp, and 100 bp to generate mappability tracks for unmodified and bisulfite-converted genomes of human (GRCh37 and GRCh38) and mouse (GRCm37 and GRCm38). We store uniquely mappable regions of these genomes in bigBed format as a track hub that can be loaded to UCSC or Ensembl genome browsers. The track hub contains one supertrack for Umap and one supertrack for Bismap. The track hub is available at http://bismap.hoffmanlab.org.

## 3 Results

### 3.1 Mappability of ENCODE ChIP-seq peaks

ChIP-seq identifies proteins present in chromatin at particular loci and often involves short-read sequencing. The ENCODE Project^5^ has performed around 1200 ChIP-seq assays on approximately 200 chromatin binding factors in more than 60 different human cell types. To show how mappability affects downstream analysis of experiments such as ChIP-seq, we quantified the mappability of narrow peaks identified in ENCODE ChIP-seq experiments. Among 1193 experiments, most peaks map uniquely. For some experiments, however, a high number of peaks overlap with non-uniquely mappable regions. Most of these experiments correspond to ChIP-seq of histone modifications with read lengths from 24 bp to 36 bp. There are two ENCODE NRF1 ChIP-seq experiments in K562 with 36bp (ENCSR000EHH) and 100bp (ENCSR494TDU and ENCSR998AJK) read lengths. For ENCSR000EHH among the 3,994 peaks called by MACS2, 219 extend into a region that is not uniquely mappable. Although the ChIP-seq signal is completely within a uniquely mappable region, MACS2 identifies a much broader peak than is warranted (Figure 3c).

**Figure 3:**
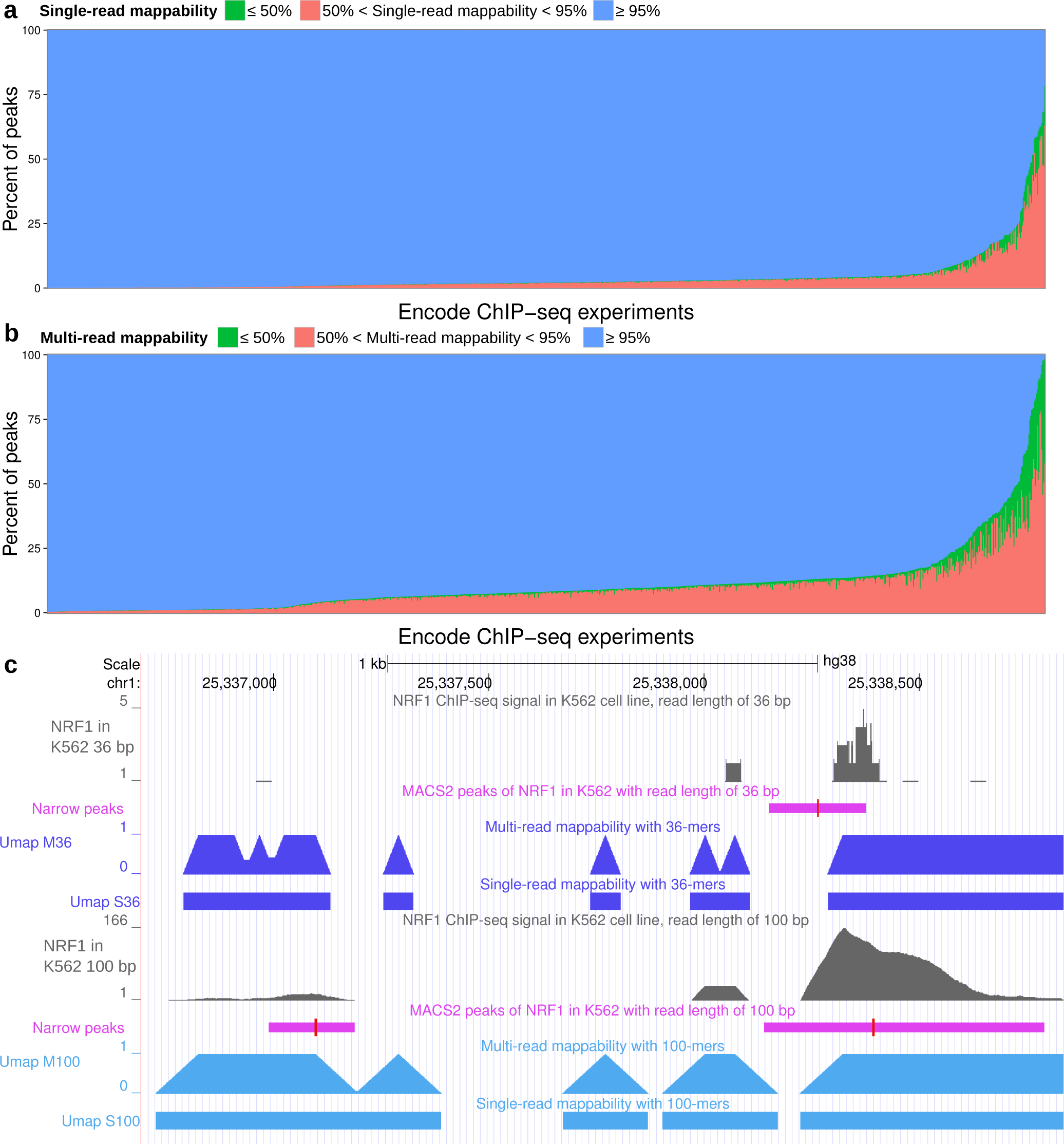
Mappability of ChIP-seq peaks in 1193 ENCODE datasets. (**a**) Single-read mappability and (**b**) multi-read mappability for narrow peaks identified in ENCODE ChIP-seq datasets. (**c**) An NRF1 narrow peak identified by MACS (purple) that is not uniquely mappable in the experiment with read length of 36 bp. The red bar in peaks indicates the summit. Signal tracks (gray) show two different replicates of this ChIP-seq experiment in K562 chronic myeloid leukemia cells (ENCODE accessions ENCSR000EHH and ENCSR494TDU, with read lengths of 36bp and 100bp respectively). Umap tracks show single-read and multi-read mappability for two different read lengths of 36 bp and 100 bp.

### 3.2 Mappability of CpG islands

CpG islands substantially overlap transcription start sites and differentially methylated regions^10^. Because CpG islands have a high number of CpGs, they are highly affected by bisulfite conversion. Thus we investigated CpG islands and the neighboring CpG shores and CpG shelves.

Even with a relatively long read length of 100 bp, 3,059/167,694 CpG annotations have zero uniquely mappable bases, as calculated by Bismap. For shorter read lengths, even more of the bisulfite-converted genome lacks unique mapping. For a read length of 100 bp, 26,510 CpG annotations are not uniquely mappable with Bismap. This represents 15.8% of all CpG annotations. The average single-read mappability of CpG annotations that are not uniquely mappable is 68.8%.

CpG islands and regions around them are often not uniquely mappable, to a lesser extent, in an unmodified genome. For example, the average single-read mappability of 15,776 CpG annotations that are not uniquely mappable in the unmodified genome is 60% with a read length of 100 bp. This is substantially lower than the average single-read mappability of the genome (92%). Also, there are 631 CpG islands that have some overlap with uniquely mappable regions of the unmodified genome, but are not uniquely mappable in the bisulfite-converted genome.

The difference in genomic mappability and CpG island annotation mappability is even more extensive for shorter read lengths. For example, for a read length of 24 bp, more than 96.84% of CpG island annotations are not uniquely mappable, but the percent of the genome that is not uniquely mappable is only 30% (Figure 4).

**Figure 4:**
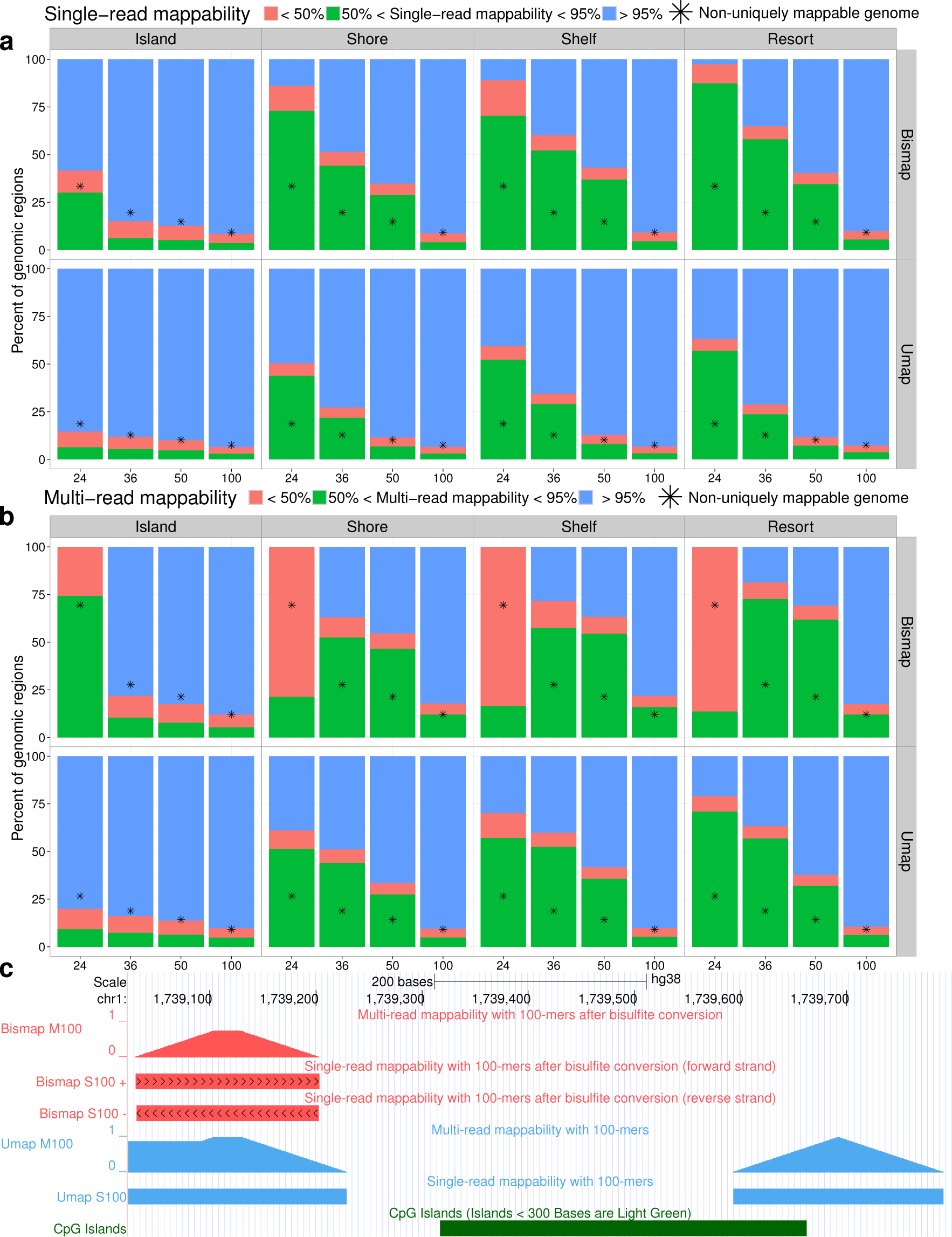
Mappability of the CpG island annotations. (**a**) Single-read mappability and (**b**) multiread mappability of CpG islands, CpG shores, CpG shelfs, and CpG resorts for a variety of read lengths. For comparison, asterisks indicate the average mappability of the whole genome at each read length. (**c**) A CpG island that is not uniquely mappable with a read length of 100 bp by Umap and Bismap. In Bismap single-read mappability tracks, chevrons pointing right indicate mappability of the + strand and chevrons pointing left indicate mappability of - strand. Multi-read mappability is calculated bases on reads that are uniquely mappable on both + strand and - strand.

**Figure 5:**
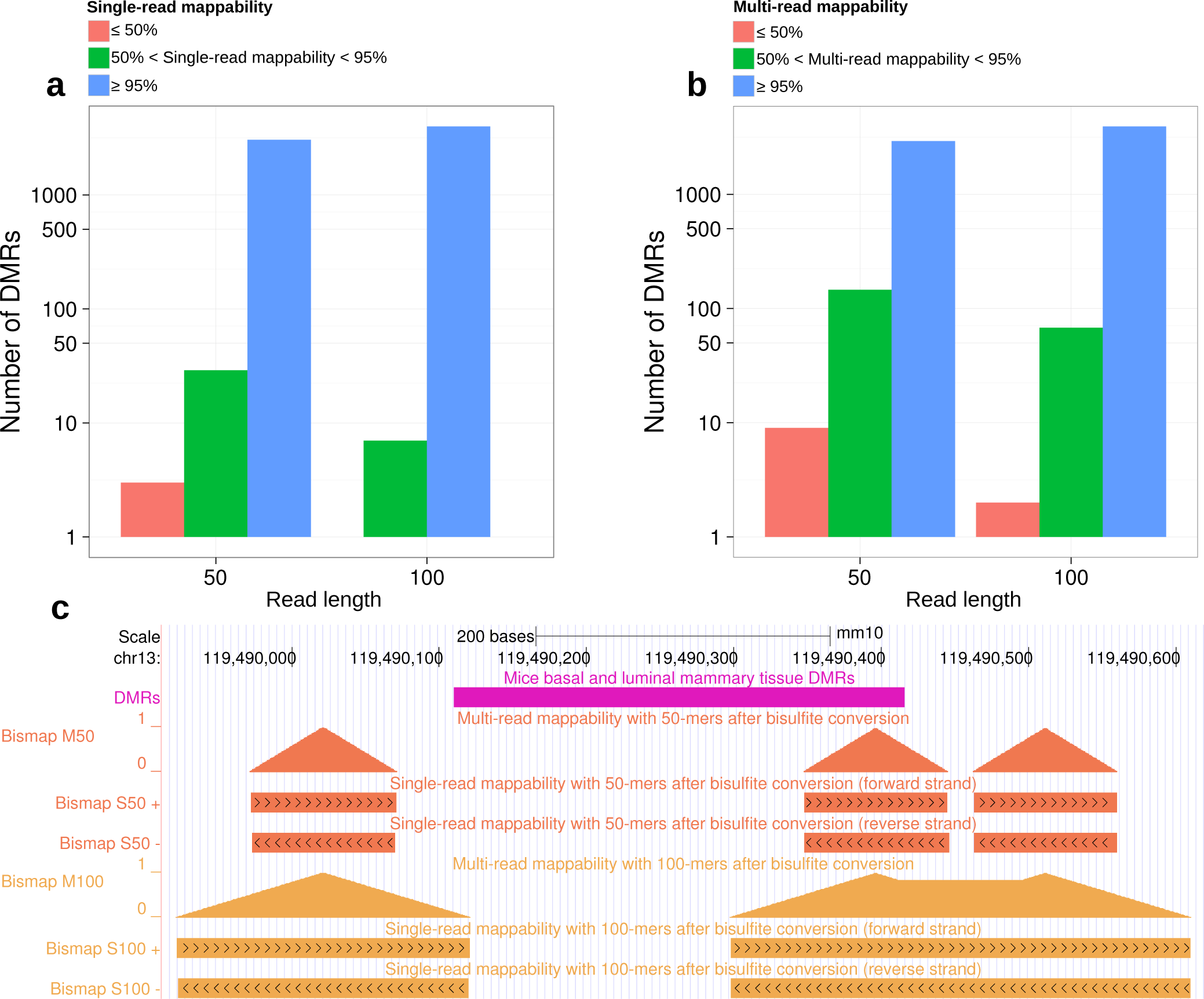
Mappability of differentially methylated regions of mice mammary basal and luminal alveolar tissues. (**a**) Single-read and (**b**) multi-read mappability of differentially methylated regions. (**c**) Example of a differentially methylated region identified with 50-nucleotide sequencing reads that is not uniquely mappable.

### 3.3 Mappability of differentially methylated regions

Many studies measure differences in methylation associated with a disease phenotype. These studies test whether each CpG’s methylation status correlates with the phenotype. Collective difference of CpG dinucleotides in a given region, however, may provide higher statistical power in assessing the association of methylation profile with disease states^21^. Cluster of CpG dinucleotides are also a more predictive feature of disease states than differences in individual CpGs^21^. BSmooth^16^ is one of the tools that identifies differentially methylated regions by estimating a smoothed methylation profile.

We compared differences in CpG methylation of basal and luminal alveolar murine mammary tissues ^13^ using BSmooth^16^. Out of a total of 965,181 CpG dinucleotides sequenced with a read length of 50bp (see Methods), 4,091 of them are not uniquely mappable. For a read length of 100bp, out of a total of 1,136,993 CpG dinucleotides, 1,980 are not uniquely mappable. For the same experimental setup, BSmooth identified 3082 differentially methylated regions for a read length of 50 bp and 3990 regions for a read length of 100 bp. For a read length of 100 bp, 17 differentially methylated regions were not uniquely mappable (single-read mappability < 100%), while for a read length of 50 bp, 8 differentially methylated regions were not uniquely mappable. This is a proof of principle that differential methylation analysis can identify false signals that are not even uniquely mappable.

DiseaseMeth^17^ catalogs publicly available methylome datasets, including 12,073 using array technologies. The cost-efficiency of these approaches has driven wide adoption. Many of these datasets, however, include probes with low mappability in the bisulfite-converted genome. The widely used Illumina Infinium methylation arrays use 50 bp probes capturing certain CpG dinucleotides^22^. Out of the 27,578 probes in the Illumina Infinium HumanMethylation27 (27K) BeadChip, 377 do not map uniquely to GRCh37, and 115 do not map uniquely after bisulfite conversion. Additionally, 304 uniquely mappable probes have low multi-read mappability, meaning that single nucleotide polymorphisms or mutations can result in probe multi-mapping (Figure 6a). Similarly, out of 485,512 probes in the Illumina Infinium HumanMethylation450 (450K) BeadChip, 84 are not uniquely mappable to GRCh37, 4,146 are not uniquely mappable after bisulfite conversion, and another 12,744 uniquely mappable probes have low multi-read mappability (Figure 6b).

**Figure 6:**
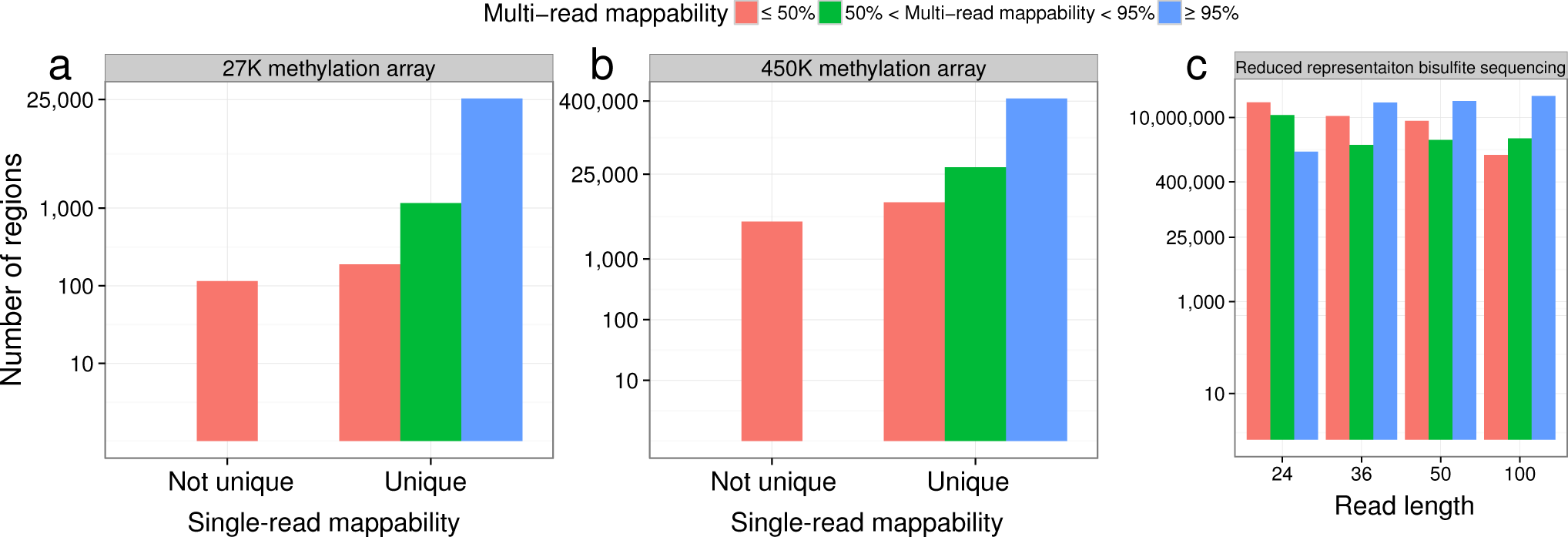
Mappability of targeted methylation assays. Multi-read mappability of probes in (**a**) the Illumina Infinium HumanMethylation27 (27K) BeadChip and (**b**) the Illumina Infinium HumanMethylation450 (450K) BeadChip. (**c**) Multi-read mappability of CpG dinucleotides found in DiseaseMeth RRBS datasets.

In addition, many publicly available RRBS datasets exist. In RRBS, only DNA fragments between 40 bp and 220 bp are selected. The majority of selected fragments, however, are approximately 50 bp ^23^. Even with a read length of 100 bp, 408,384 (1.18%) of CpG dinucleotides in RRBS experiments of DiseaseMeth database did not map uniquely (Figure 6c).

## 4 Discussion

### 4.1 The importance of considering mappability in analysis

In several examples we showed how mappability must be considered in analysis of sequencing data. One needs to examine, however, the extent of genomic variation which affects mappability calculations. Genetic variants specific to each sample make it impossible to know the exact mappability. We introduced a measure called multi-read mappability for addressing this issue. Genomic regions with higher multi-read mappability are less prone to be biased by genetic variants and sequencing errors.

In ENCODE ChIP-seq experiments using short read lengths, we found many examples where signal was within a uniquely mappable region but peaks identified by peak caller had substantial overlap with non-uniquely mappable regions. More than 50% of ChIP-seq data in the ENCODE Data Coordination Center use reads shorter than 36 bp. Consortia such as ENCODE and Roadmap have spent hundreds of millions of dollars to perform these experiments, which they won’t repeat any time soon. This shows the importance of using the mappability information to analyze sequencing data, especially when the read length is short. In fact, we initially developed Umap as part of the ENCODE uniform analysis pipeline^5^ to avoid such problems.

In Bismap, we convert all cytosines to thymines in the forward strand, and all guanines to adenines on reverse strand, just as alignment algorithms such as Bismark^3^ or BWA-meth^4^ do. In practice, chemical resistance or sample-specific genetic variation may retard bisulfite conversion. This makes it impossible to estimate the exact mappability for a bisulfite converted sample. When performing bisulfite sequencing on different mouse strains, using the same reference genome for each introduces massive bias in bisulfite sequencing data analysis^24^. Ideally, one would align data from each strain to a reference genome specific to that strain. When one lacks a strain-specific reference genome, Bismap at least allows us to quantify how and where genetic variation affects reliability of bisulfite sequencing results. While Bismap assumes complete bisulfite-conversion, Umap assumes none. By comparing the results of the two methods, we can understand the range of bisulfite-conversion effects on mappability.

While paired-end sequencing with lengths greater than 100 bp has become more common, most publicly available datasets such as ENCODE have used shorter reads. Out of 3,483 ENCODE ChIP-seq experiments, 3,033 use single-ended sequencing, and 2,228 have read lengths of 36 bp or shorter. Out of the 142 ENCODE RRBS datasets, 140 (98.6%) have a read length of 36 bp or shorter. In addition, commonly used array technologies such as the 450K array uses 50 bp probes and multi-read mappability of some of the probes is low. This allows multi-mapping due to genetic variation and decreases data quality in these regions as it has been noted before^25^. Although only a small fraction of all probes do not map uniquely (1.8% in the 27K array and 0.87% in the 450K array), one must still use caution when interpreting methylation signal—or the lack thereof—in these regions. In fact, multi-mapping probes have lead to false discovery of autosomal sex-associated DNA methylation in at least one study^26^.

In our analysis of whole genome bisulfite sequencing data of mouse mammary tissue, ~0.1% of CpG dinucleotides were not uniquely mappable with 50 bp reads. We removed reads with a mapping quality of less than 10 and only counted CpG dinucleotides that had a minimum coverage of 3 reads in all of the 5 different whole genome bisulfite sequencing datasets. Given this stringent filtering, the chance of observing any non-uniquely mappable read is 10^−15^ which is much less than our observation (0.1%). Such CpG dinucleotides must be excluded from analysis. RRBS usually involves filtering fragments to only include those that are 40 bp-220 bp, and most RRBS reads are 50 bp or less^23^. This causes a major issue for mapping of these reads.

In paired-end sequencing, short regions from both ends of a longer fragment are sequenced. This provides a long read more likely to map uniquely to the genome. The length of these fragments varies considerably in size. One can still use Umap or Bismap to identify the mappability for a range of k-mers that represent the variation in fragment length of any given sequencing library.

In RNA-seq, gap alignment algorithms account for splicing. Different software and user defined parameters handle multi-mapping reads differently which can be a source of error. Robert and Watson^27^ recommend to assign multi-mapped reads to a group of genes instead of removing them. They show that this approach accurately recovers a significant portion of the data.

### 4.2 Other methods for mappability

Bias Elimination Algorithm for Deep Sequencing (BEADS^28^) also defines a mappability measure that is obtained by identifying uniquely mappable 35-mers of the genome. Based on the assumption that each read identifies a longer 200-mer, BEADS extends uniquely mappable 35-mers to 200 bp, and calculates the fraction of reads that span a given genomic position. BEADS uses a cutoff of 25% mappability to filter signals that might bias a study. Extending the 35-mer mappability to 200 bp, however, defines the exact mappability for neither 35-mers nor 200-mers.

PeakSeq^29^, uses an algorithm similar to Umap and identifies the single-read mappability in 1kbp windows of the genome. PeakSeq filters out ChIP-seq signals with low mappability in each window by comparing it to a simulated background of reads with Poisson distribution.

Model-based one and two Sample Analysis and inference for ChIP-Seq Data (MOSAiCS)^30^ uses a mappability measure similar to multi-read mappability for preprocessing of data. While Umap’s multi-read mappability calculates the percent of uniquely mappable *k*-mers that span each nucleotide, MOSAiCS calculates the percent of *extended uniquely mappable k-mers* for calculating its mappability score. In comparison to other mappability measures, Umap’s multi-read mappability has the advantages of specificity to an exact read length and efficient calculation for any read length.

## Acknowledgements

We would like to thank Scott M. Lundberg for providing us with GRCh38-aligned BAM files of the ENCODE ChIP-seq datasets. We also thank Carl Virtanen and Zhibin Lu at the University Health Network High Performance Computing Centre and Bioinformatics Core for technical assistance. This work was supported by the Canadian Cancer Society (703827 to M.M.H.), Ontario Institute for Cancer Research (OICR), the Natural Sciences and Engineering Research Council of Canada (RGPIN-2015-03948 to M.M.H. and RGPIN-435512-2013 to C.E.), the University of Toronto McLaughlin Centre (MC-2015-16 to M.M.H.), and the Princess Margaret Cancer Foundation.

## Competing interests

The authors declare that they have no competing interests.

